# Real-time Taxonomic Characterization of Long-read Mixed-species Sequencing Samples in Sorted Motif Distance Space: *Voyager*

**DOI:** 10.1101/2024.04.13.589333

**Authors:** Sverre Branders, Manfred G. Grabherr, Rafi Ahmad

## Abstract

Recent advances in long-read sequencing technology enable its use in potentially life-saving applications for rapid clinical diagnostics and epidemiological monitoring. To take advantage of these enabling characteristics, we present *Voyager*, a novel algorithm that complements real-time sequencing by rapidly and efficiently mapping long sequencing reads with insertion- and deletion errors to a large set of reference genomes. The concept of *Sorted Motif Distance Space* (*SMDS*), i.e., distances between exact matches of short motifs sorted by rank, represents sequences and sequence complementarity in a highly compressed form and is thus computationally efficient while enabling strain-level discrimination. In addition, *Voyager* applies a deconvolution algorithm rather than reducing taxonomic resolution if sequences of closely related organisms cannot be discerned by *SMDS* alone. Using relevant real-world data, we evaluated *Voyager* against the current best taxonomic classification methods (Kraken 2 and Centrifuge). *Voyager* was on average more than twice as fast as the current fastest method and obtained on average over 40% higher species level accuracy while maintaining lower memory usage than both other methods.

## Introduction

The journey to the next generation of medical care in the face of infection was headed off when Oxford Nanopore Technology (ONT) first introduced true real-time sequencing technology. Combined with its portability, low cost, and ease of library preparation, a path to the use of sequencing technology in routine diagnosis has become accessible. To arrive at such new rapid clinical diagnostics, novel computational methods are needed to characterize infectious pathogens quickly and accurately by taking advantage of these real-time capabilities. In a clinical context, if patient samples obtained from body fluids, e.g., blood sputum, urine, etc., can be quickly analyzed to characterize pathogens, this will lead to faster and informed treatment, reduced patient time in the hospital, and improved patient outcome^1^. Likewise, fast and user-friendly characterization of infections in the veterinary and livestock sector can relieve the need for preventative antibiotics and prevent the spread of infection to the rest of the herd^2^. Advances in real-time sequencing technologies have the potential to enable such rapid diagnosis in clinical microbiology. However, the process needs to be complemented by fast computational analyses to deliver results in true real-time^1^.

Previously, the problem of identifying species in a mixed-species sequencing sample has been addressed by either mapping the reads to a set of known reference genomes^3,4^ or by analyzing the local nucleotide characteristics against a reference database^5^. Either case relies on matching up short consecutive strings of nucleotides, e.g., k-mers, allowing for fast look-up or analyzing k-mer frequencies, compensating for isolated sequencing errors in between perfect matches. While these techniques are well-suited for short read sequencing methods, such as Illumina, where errors manifest themselves predominantly as single nucleotide substitutions, they do not work optimally for long reads with insertion/deletion error patterns^6^. Initially, ONT sequencing had an error rate of over 10%, with most errors caused by insertions or deletions^7^. In recent years, the error rate has steadily decreased. However, most remaining errors are still caused by frequent short deletions or, less frequently, long insertions^8^.

Here, we present *Voyager*, a new method that maps reads in *Sorted Motif Distance Space* (*SMDS*) and infers the taxonomic composition of a sample from these mappings across a complete sequencing run.

### Novelty

*Voyager* works by measuring the distances between exact matches of short motifs, implementing a concept that borrows from optical restriction map alignments^9^. Unlike k-mers, i.e., local short consecutive sequences, these distances span dozens, hundreds, or even thousands of nucleotides, depending on the choice of the motif. When combining consecutive distances *into local distance vectors* of a certain fixed *window size*, these vectors are highly specific to the sequences they originate from since the exact distances are unlikely to occur by chance and in the same order in a different genome.

To map error-containing sequencing reads, instead of using distance vectors directly, *Voyager* sorts these vectors into *vectors of ranks*, effectively making the ranks the key for look-up: the ranking is preserved in the presence of insertions and deletions, so long as the sorted order is not changed. Given a fixed window size *n*, the number of possible permutations is *n* factorial, e.g., for *n*=14, there are 8.71×10^10^ permutations. When using a motif that occurs every 100 nucleotides on average, this *permutation space* is more than three orders of magnitudes larger than the number of *vectors of ranks* found in a mammalian genome and still two orders of magnitudes above large plant genomes.

As a result, *Voyager*’s algorithm (a) is computationally efficient due to a sparse and long-range search space; (b) compresses the search index, thus memory usage, by omitting the sequences in between motifs and only using the distances; and (c) provides high specificity, to the degree that individual strains are discernable: instead of going up in taxonomy if strains, species, clades, or taxa are not discernable, *Voyager* applies a deconvolution step, where the underlying composition of a sample is estimated using the non-negative-least-squares (NNLS) algorithm^10^.

In the following, we describe the algorithms in detail, explain our choice of parameters, and then demonstrate how this conceptually simple but smart algorithm compares against existing methods.

## Results

### *SMDS* provides an efficient and specific mapping algorithm

The working mechanism of *Voyager*’s *SMDS* mapping algorithm is illustrated in Figure 1. The Figure shows a 274 bp section of an alignment of a query sequence containing six motif matches to an ONT sequencing read of the corresponding gene region. In nucleotide space, the alignment includes 23 INDEL mismatches and eight substitution mismatches. Due to substitution errors, the query sequence contains an additional two motif matches. Despite the high INDEL errors in the ONT read sequence, (given a window size of four), two of *the ranked distance vectors* are perfectly conserved. Where errors affect motif matches, the sorted order changes in the windows containing these, and they thus no longer match. This mechanism gives *Voyager* its discriminative ability.

**Figure 1:**
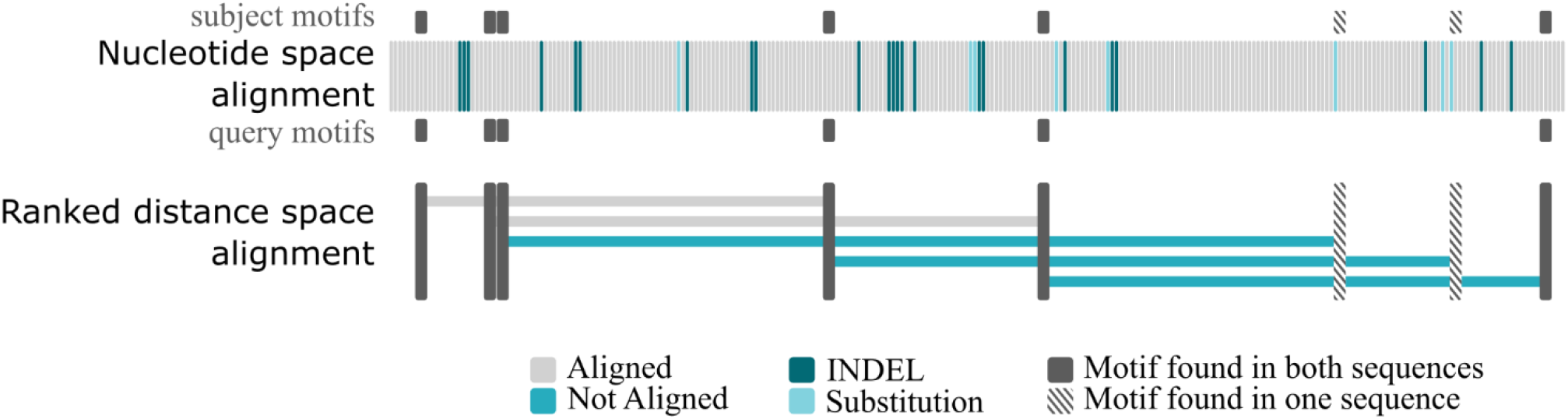
Illustration of the *Voyager ranked distance space* matching algorithm. The figure shows an alignment of a sequence to an error containing ONT read of the corresponding gene, in nucleotide space and in *Sorted Motif Distance Space* (*window size* = 4).

Moreover, the *ranked distance space* conserves the similarity between the two sequences while greatly compressing the search space. When constructing *local distance vectors* of the NCBI RefSeq^11^ prokaryotic Representative Genomes using the soft-mer motif “AST” (where “S” is either C or G), on average, the distance between motifs is 106 nucleotides (σ 328) with 95% of the distances falling between 30 nt and 329 nt (Figure 2a). The average distance between motifs “CWG” (“W” is A or T) (Figure 2b) is 38 nt (σ 74), with 95% of distances between 19 nt and 86 nt. When using motif “AGCT” (Figure 2c), the mean distance increases to 658 nt (σ 868) with 95% of distances above 183 nt and below 3374 nt. Motif “CASTG” (Figure 2d) results in a mean distance of 1407 nt (σ 1259) with 95% in the range of 393 nt to 5088 nt.

**Figure 2:**
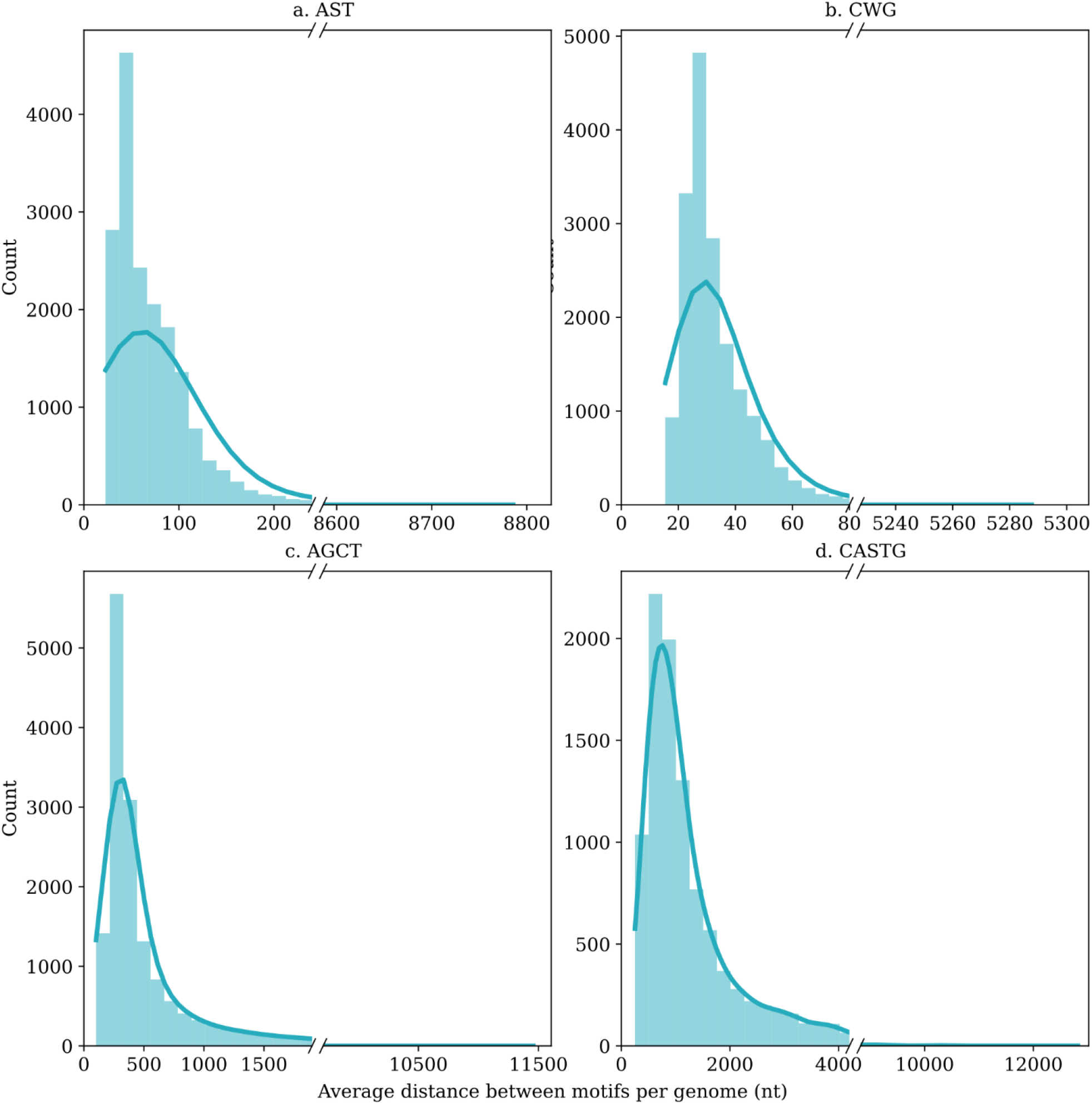
Histograms of the average distances between motif matches. The histograms show the average distance in nucleotides between exact motif matches for all genomes in the NCBI RefSeq prokaryotic Representative Genomes collection. The distances were obtained using different motifs: (a) AST, (b) CWG, (c) AGCT, and (d) CASTG.

### Voyager achieves Strain level Specificity

We compared the specificity of *Voyager*’s algorithm between real sequence data and the theoretical optimum, i.e., reads simulated from the genomes directly and without any sequencing errors. We first extracted 100 randomly chosen *vectors of ranks, window size* 14, motif “AST”, per permutation from the *Escherichia coli* O25:H4 genome (NCBI GenBank assembly (<u>GCA_030507855.1</u>), and mapped these into sixteen bacterial genomes. Figure 3a shows the distribution of the hits, where *E. coli* O25:H4, as expected, receives the highest number of hits. Two other strains of *E. coli*, K12 and Sakai, also match the reads, albeit at a much lower hit abundance, which we attribute to the high sequence similarity between the strains. However, with increasing taxonomic distance, the hits approach zero. We then repeated the analysis for real sequencing data from a sample obtained from strain *E. coli* O25:H4 (ENA accession <u>ERR11001111</u>). Figure 3b shows that the experiment performed on real sequencing data results in very similar distributions, with distributions only slightly skewed towards more false positives, which can likely mostly be attributed to sequencing errors affecting the motifs sequences and/or the ranked order of distances between them.

**Figure 3:**
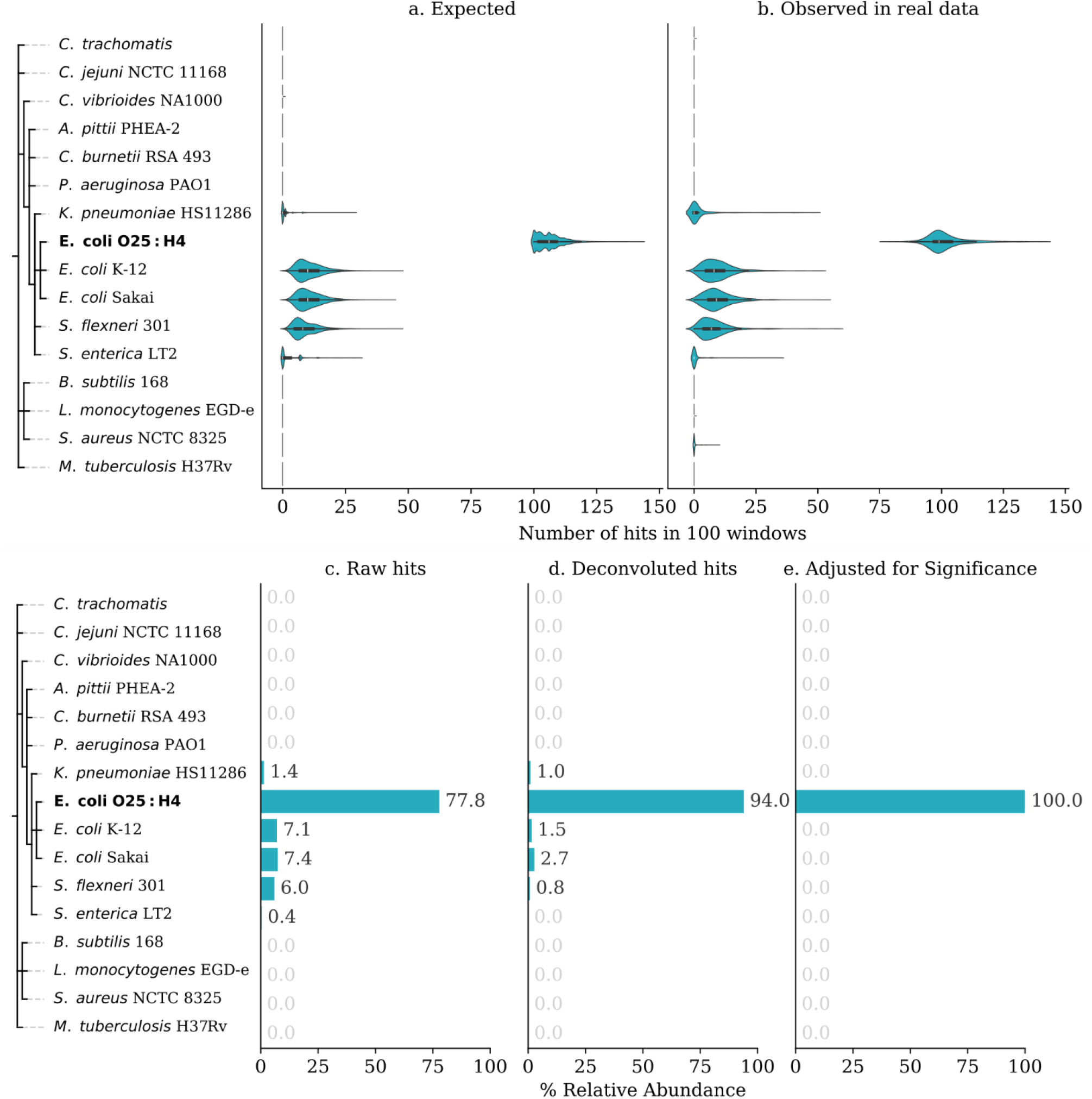
Specificity of *Voyager*’s mapping algorithm in *permutation space* and against real data. (a) Expected specificity of the *Voyager* mapping algorithm measured through random permutations of 100 randomly chosen *vectors of ranks, window size* 14, motif “AST” (b) Observed specificity in real sequencing data using random permutations in the same manner. (c) Raw hits obtained after mapping a real sequencing sample using the *Voyager* algorithm. (d) Hits obtained from the same sample after deconvolution was applied. (e) Hits reported after adjusting for significance.

In quantifying specificity, the raw hits obtained from the mapping algorithm (Figure 3c) clearly follow the expected distributions, providing a strain level specificity of 77.8%, where most false matches occur in closely related strains. After *Voyager*’s deconvolution (Figure 3d), these false matches can be mostly corrected, obtaining a 94% strain-level sensitivity. The deconvolution algorithm cannot account for individual variation in error rate and bias between sequencing runs, which is likely why it cannot correct for 100% of false hits. Finally, *Voyager* estimates the statistical significance of each genome. When correcting for this significance, *Voyager* obtained strain-level discrimination with 100% sensitivity in this case (Figure 3e).

We compared *Voyager*’s performance to the most popular and performant current methods. Centrifuge^3^ and Kraken2^5^ are widely used, with citations of nearly 1000 and over 2100 (> 5000, including related publications), respectively. Furthermore, their performance has repeatedly proven reliable with ONT platforms, and ONT has integrated them into their pipelines^12,13^. Notably, both tools’ results can be directly compared to *Voyager*’s, and the same databases can be used. In Figure 4, the specificity of *Voyager*’s algorithm is compared directly to Centrifuge and Kraken2. While each tool obtains similar species level accuracy, false hits obtained by *Voyager* occur in closely related species. In contrast, false hits obtained by Centrifuge and Kraken2 occur in a wider taxonomic spread, suggesting that *Voyager* is less prone to random noise. This characteristic may result in comparatively fewer false positive identifications in real-world data by *Voyager*.

**Figure 4:**
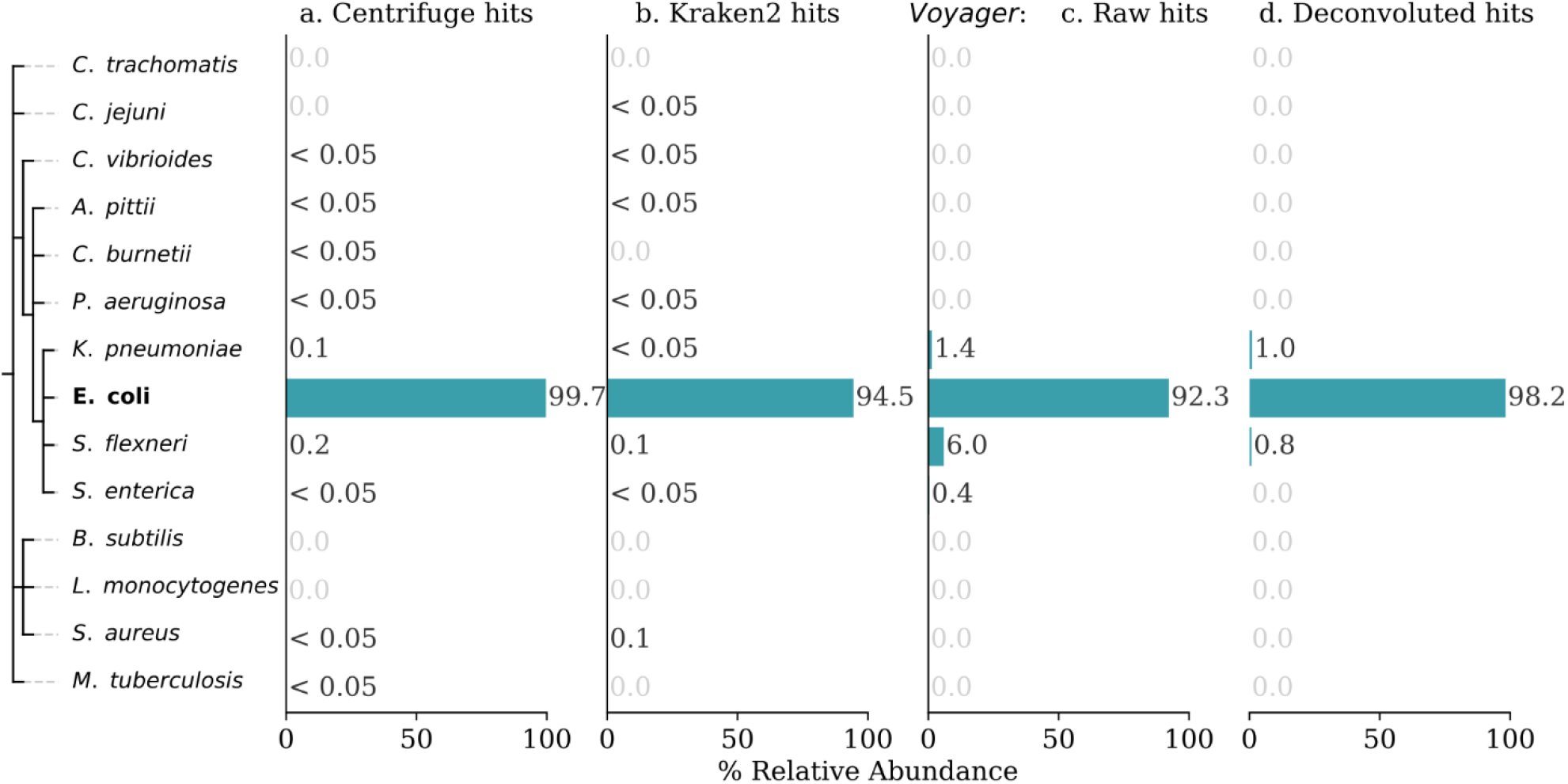
Specificity of *Voyager*’s mapping algorithm compared to the specificity of Centrifuge and Kraken2 on species level. (a) Relative abundance of reads mapped to each species using Centrifuge. (b) Relative abundance of reads from the same sample mapped to each species using Kraken2. (c) Raw hits obtained from the *Voyager* mapping algorithm on the same sample. (d) Hits reported by *Voyager* after applying deconvolution.

### Higher Speed and low Memory Footprint

We collected datasets from all available ONT sequencing platforms, flowcells, and relevant sample types to establish expected performance in clinical, animal, and limited environmental community samples, including blood infection data obtained using MinION^14^ (ENA <u>PRJEB60525</u>) and Flongle^15^ (ENA <u>PRJEB49072</u>), urine infection data^16^ using MinION (ENA <u>PRJEB73819</u>) milk mastitis data^2^ using MinION (ENA <u>PRJEB53532</u>), and environmental mock community data^17^ using GridION and PromethION (ENA PRJEB29504). Sample files containing low amounts of data (< 100,000 reads) were excluded. To compare *Voyager*’s speed with current methods, these sequencing files (n = 51) were mapped to the NCBI RefSeq prokaryotic Reference Genomes collection using Centrifuge, Kraken2, and *Voyager*. CPU times were measured and compared. On average, *Voyager* used 3.53X less CPU time than Centrifuge and 2.29X less CPU time than Kraken2 (Figure 5a). Differences in CPU time with Centrifuge varied greatly (σ 2.07) where *Voyager* was at most 11.36X faster and at the lowest 1.80X faster than Centrifuge. Differences in CPU time with Kraken2 fell within 1.71X and 2.73X (σ 0.28).

**Figure 5:**
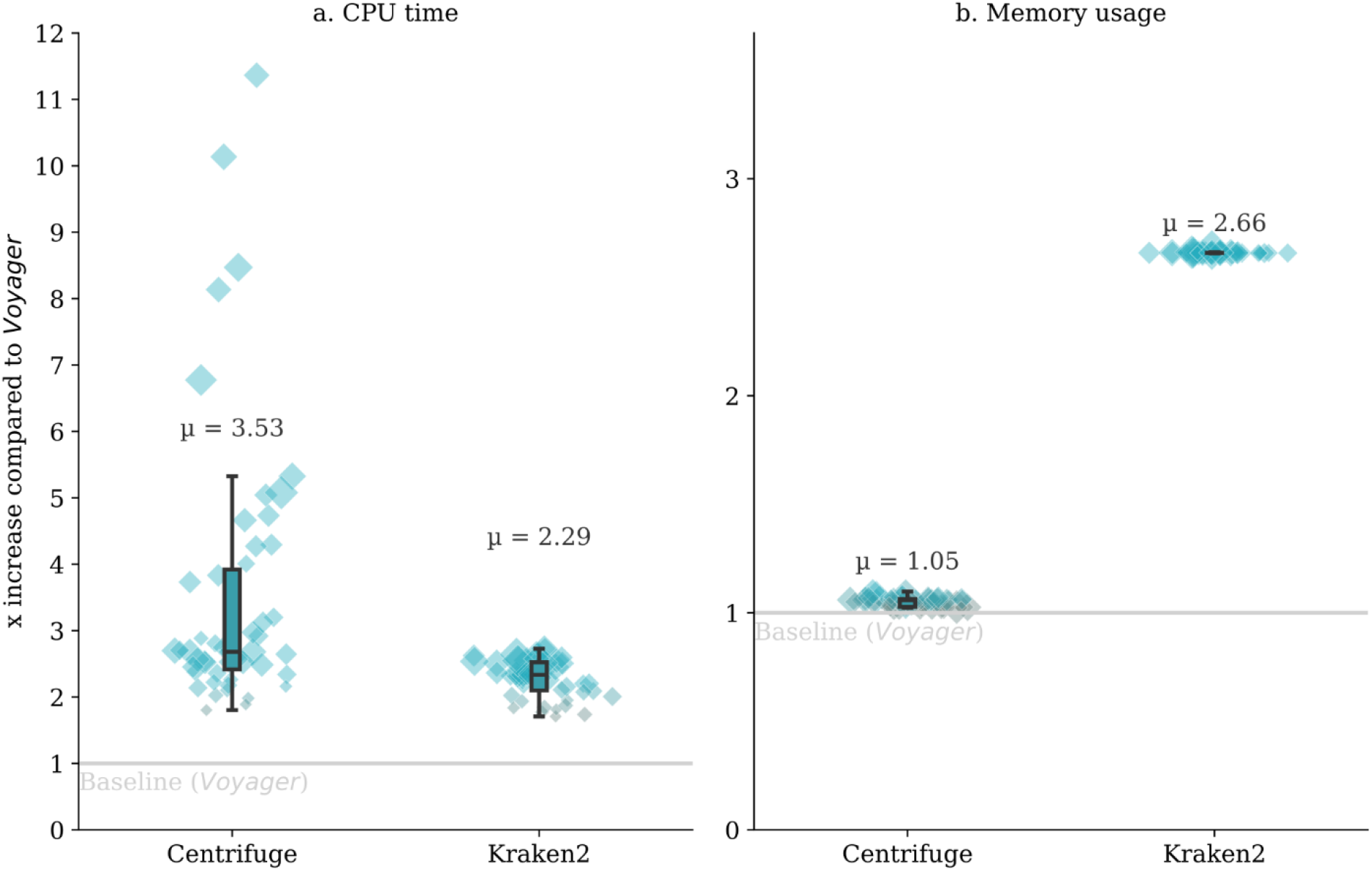
CPU Time and Memory usage of Centrifuge and Kraken2 compared to *Voyager*. Used CPU time and peak Random Access Memory (RAM) X increase of tools compared to *Voyager* for files containing more than 100,000 reads. The size of the point reflects the relative size in KB of the sequencing sample. (a) CPU time increase for each sample compared to *Voyager* using the same database for Centrifuge and Kraken2. (b) Peak RAM usage X increase for each sample compared to *Voyager* using the same database for Centrifuge and Kraken2.

To investigate memory usage, indexes were created using the NCBI RefSeq prokaryotic Representative Genomes collection (currently including 18 342 assemblies) for Centrifuge, Kraken2, and *Voyager*. Centrifuge’s index used roughly 48 GB of Random Access Memory (RAM), Kraken2’s index used ca. 121 GB, and *Voyager*’s ca. 45 GB. The same sequencing files (n = 49 due to missing data) were mapped using these indexes and the difference in peak RAM usage was plotted (Figure 5b). As expected, memory usage varied little, as most memory usage for each tool is attributable to the index size. Centrifuge’s memory usage was comparable to *Voyager*’s, using, on average, 1.05X as much RAM (σ 0.02). Kraken2 used, on average, 2.66X as much RAM as *Voyager* (σ 0.002).

### Voyager Outperforms other tools at Strain-level Classification of Real-world Data

We constructed a database containing all bacterial and fungal species in the test data set (n = 52) and compared the ability to accurately detect species between Centrifuge, Kraken2, and *Voyager*. The percentage relative abundance of true positive hits was calculated for each tool based on the ground truth for each tested sample. For *Voyager*, this was done with the raw hits obtained from the mapping algorithm (Figure 6a) as well as with the deconvoluted hits (Figure 6b). In 26 of the 52 tested samples (50%), deconvolution improved *Voyager*’s true positive hit abundance from an average of 89.7% to 93.0% with a mean increase of 3.30% (σ 0.0373) over the raw hits. For these 26 samples, Centrifuge and Kraken2 had a true positive hit abundance of 50.5% and 45.7%, respectively. In 13 samples (25%), *Voyager*’s raw hits were sufficient for 100% accurate species-level identification. The average true positive abundance for these 13 samples was 37.6% and 21.1% for Centrifuge and Kraken2 respectively. For 12 of the tested samples (23.1%), the deconvoluted true positive hit abundance (μ 53.2%) was marginally lower than the raw hit true positive abundance (μ 52.6%) with a mean decrease in true positive abundance of 0.58% (σ 0.0038). The mean true positive hit abundance for these twelve samples was 18.1% and 40.5% for Centrifuge and Kraken2 respectively.

**Figure 6:**
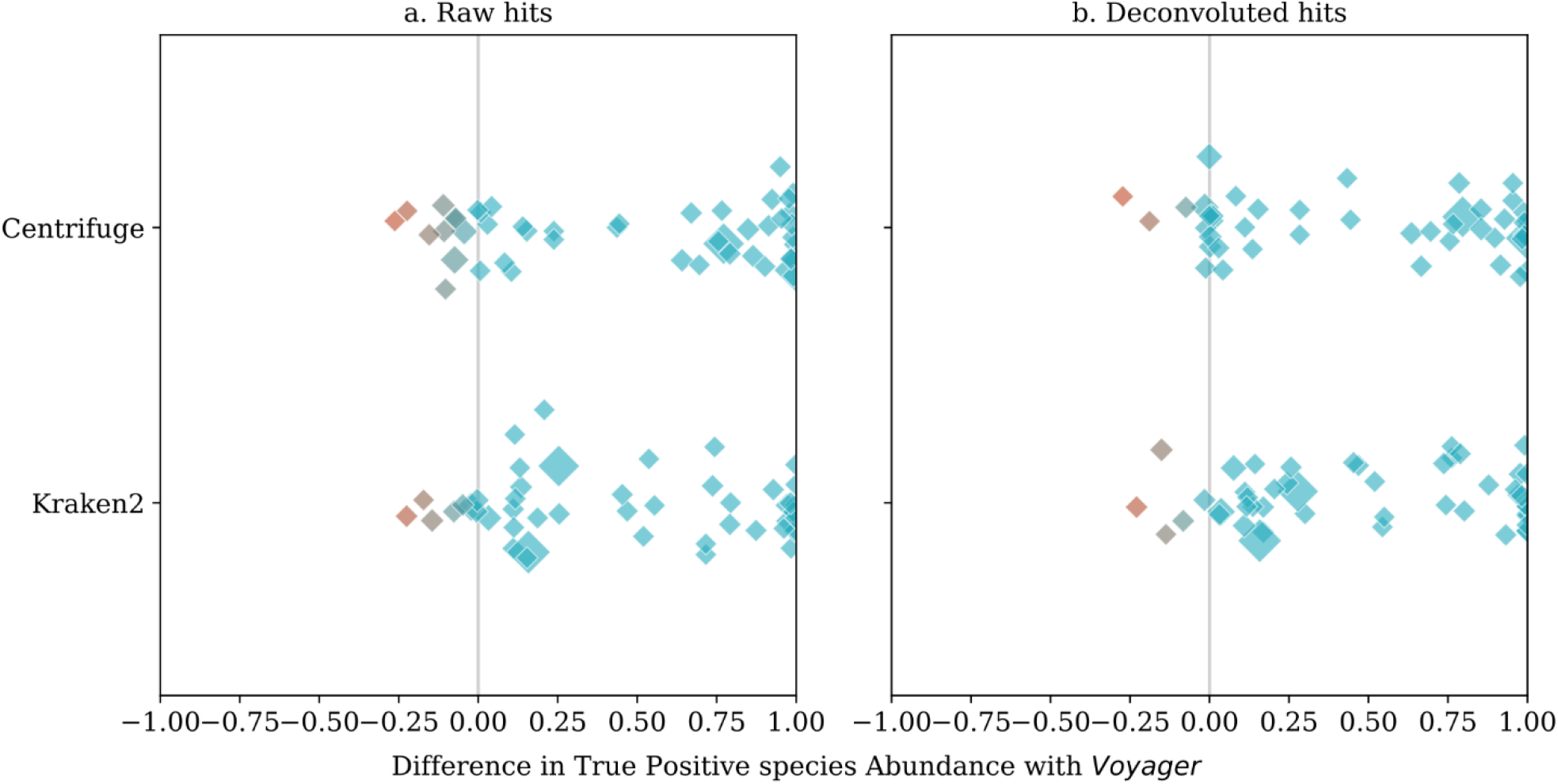
Difference in true positive species abundance of Centrifuge and Kraken2 compared to *Voyager*. The size of the point reflects the relative size in KB of the sequencing sample. (a) Difference in true positive hit abundance at the species level (according to known ground truth for each sample) with *Voyager*’s raw hits for Centrifuge and Kraken2. (b) Equivalent difference in true positive hit abundance with *Voyager*’s hits after deconvolution.

For 37 of the samples (71.2%), *Voyager* outperformed both Centrifuge and Kraken2 in terms of species level accuracy. In these samples, Centrifuge had an average of 33.0% true positive hit abundance, Kraken2 an average of 27.0% and *Voyager* an average of 91.0%. In ten samples (19.2%), Centrifuge had higher species-level accuracy (μ 74.8%) than *Voyager* (μ 68.7%) with a mean difference of 6.0% (σ 0.0944). Kraken2 had higher species-level accuracy in seven samples (13.5%), with a mean true positive hit abundance of 62.3% compared to 53.5% for *Voyager*, with a mean difference of 8.8% (σ 0.0885). Overall, mean species-level accuracy was 38.6% for Centrifuge, 37.5% for Kraken2, and 83.6% for *Voyager*. On average, *Voyager*’s accuracy was 44.8% (σ 44.41) higher than Centrifuge and 46.1% (σ 41.44) higher than Kraken2.

Since (for most samples) we only have information about the presence/absence of species, we could not directly evaluate the equivalent false positive and false negative hit abundance. Therefore, we provide the following metrics instead (Figure 7); we define a true positive (TP) as a species present in the sample correctly detected (at least one hit), a false positive (FP) as a species being detected despite not being present in the sample, and a false negative (FN) as a species present in the sample not being detected. Then, the accuracy metrics are calculated as follows:

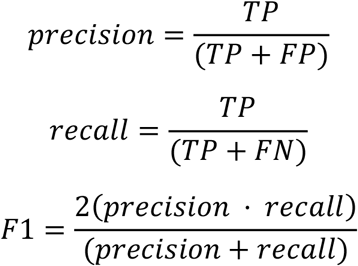

**Figure 7:**
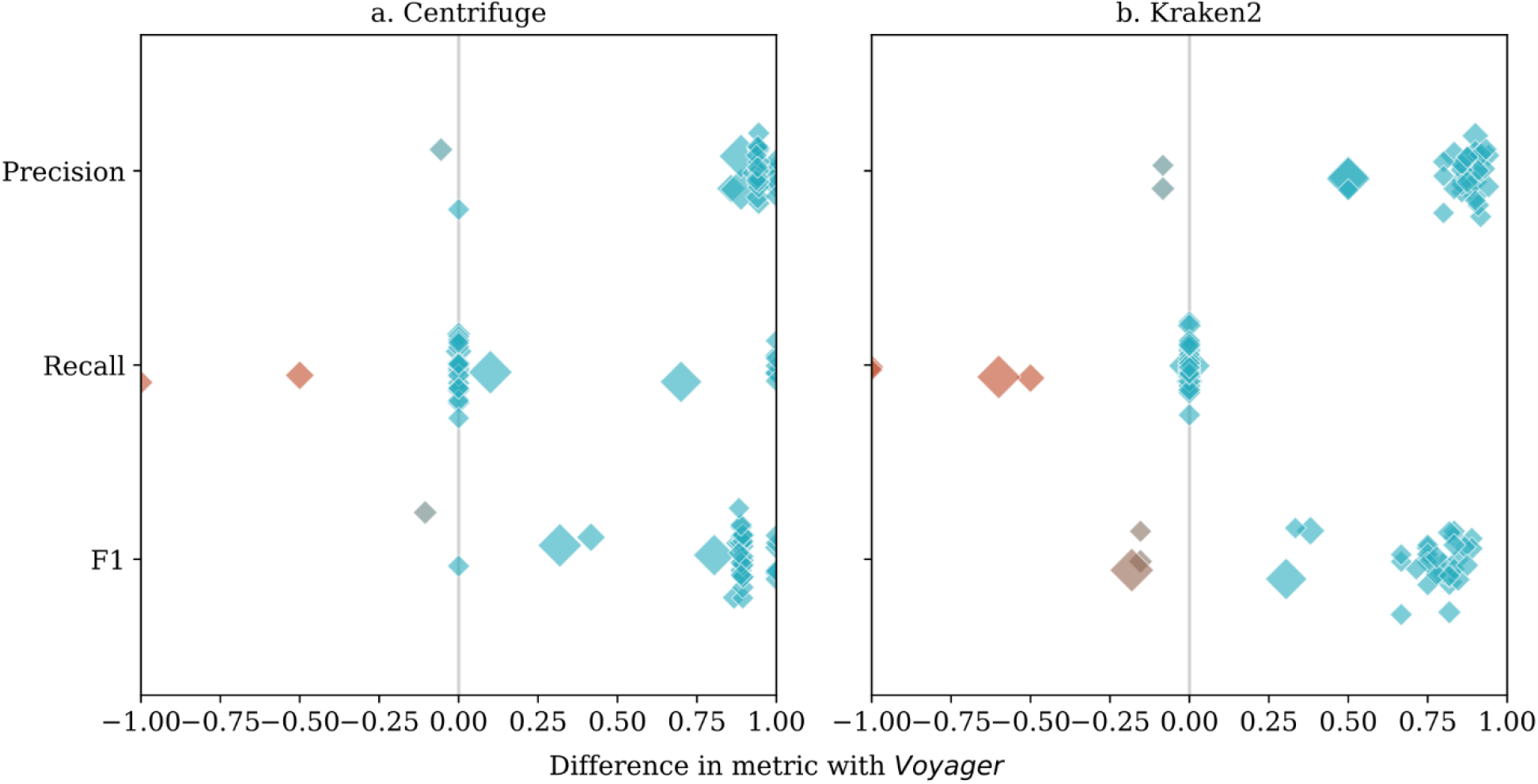
Difference in additional performance metrics of Centrifuge and Kraken2 compared to *Voyager*. Precision, recall, and F1 scores of (a) Centrifuge and (b) Kraken2 were compared to *Voyager*. The size of the point reflects the relative size in KB of the sequencing sample. True positives, false positives, and false negatives used for calculating the metrics are defined in terms of presence/absence at the species level.

In 49 out of 51 tested samples (96.1%), *Voyager* had equal or higher recall and precision than Centrifuge. *Voyager* had higher precision but lower recall than Centrifuge in one additional sample. In 48 out of 52 tested samples (92.3%), *Voyager* had equal or higher recall and precision than Kraken2, and in two additional samples, *Voyager* had higher precision but lower recall than Kraken2. Overall, *Voyager*’s F1 scores were 83.8% (σ 0.221) higher than Centrifuge’s and 67.5% (σ 0.254) higher than Kraken2’s F1 scores.

## Discussion

The ultimate goal of using sequencing technology in healthcare settings is to provide clinically relevant information at minimal turnaround time, high accuracy in clinical performance, and low cost. While advances in biotechnology have significantly sped up data acquisition, a new generation of analysis tools will enable data processing in real-time, while sequencing is ongoing and data is generated by the sequencing machine. This combination and integration allow rapid and precise diagnostics of bacterial infections, monitoring and managing disease outbreaks, and analysis of environmental samples for, e.g., surveying freshwater quality.

*Voyager*’s algorithm solves a long-standing problem in processing long read sequences, which is to rapidly and efficiently map long reads with insertion- and deletion-errors to a large set of reference genomes. Using *Sorted Motif Distance Space* allows for a high level of compression of sequencing data while maintaining specificity. This enables *Voyager to* demonstrate high speed compared to other tools while maintaining relatively low memory usage. Moreover, we have shown that using SMDS provides very high specificity and can achieve strain-level discrimination, which is vital for, e.g., distinguishing antibiotic resistant strains or epidemiological monitoring. *Voyager*’s genome deconvolution algorithm further facilitates identifying the exact strain, which largely corrects for false signals caused by sequence similarity. We have demonstrated this method’s potential on relevant real-world datasets and shown it can obtain higher species-level classification accuracy than current tools.

One possible caveat of *Voyager* is that it doesn’t find the best match on a per read basis with current ONT sequencing error rates, since errors do interfere with the motifs. Rather, *Voyager* considers the whole sequencing run to estimate the most probable taxonomic composition. However, this is merely a function of error rates, which, in the case of ONT sequencing, have been steadily declining over the last few years, and we expect the read quality to increase even further. In that regard, *Voyager*’s algorithm is “future-proof”. Lower error rates may enable changes of parameters like *window size* and motif length that could further improve *Voyager*’s efficiency. Furthermore, we expect that long read error rates will one day be low enough so that the *ranked distance vector* method can also be applied to, e.g., Whole Genome Assembly (WGA), both for single organisms and in particular for mixed-species metagenomic samples. In the meantime, taxonomic classification is a prime application for *Voyager*’s algorithm because, as we have shown, even a relatively low read pass rate still delivers accurate results.

*Voyager* is also unique because it comes with a free and easy-to-use graphical and interactive tool that allows for monitoring results and building a database of reference genomes. This intuitive interface makes *Voyager* accessible to non-experts in bioinformatics while retaining the ability to be deployed in classic bioinformatics pipelines. In addition, developers can use the API provided by *Voyager* to pass output in real-time to any application. Its goal is to provide simple yet comprehensive information about the most likely composition of the sample in true real-time. To this end, *Voyager* reports probability values, which indicate the likelihood of the sample containing that genome. In addition to this probability, *p-values* are also provided that indicate the confidence in this probability value. With a growing need for rapid diagnostics applications, this may enable metagenomic whole genome sequencing to be more accessible in, for example, clinical settings where bioinformatics expertise has often been a barrier to adoption^18^.

## Methods

### Read Mapping

We define a soft-mer motif as an IUPAC degenerate nucleotide code of short length and its reverse complement. For example, when supplying *Voyager* with the soft-mer “AMC”, *Voyager* will look for exact matches of “AAC”, “ACC”, and their reverse complements “GTT”, “GGT”, in a given sequence and record their absolute starting positions. Once the absolute positions of exact motif matches are known in a sequence, *Voyager* constructs a *local distance vector* of that sequence. Let

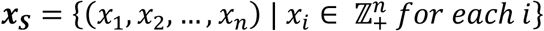

be the absolute starting positions of exact matches of a soft-mer motif *M* in a sequence *S*. Then the *local distance vector* of *S*,

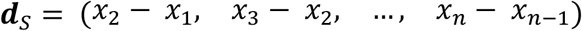

represents the local distances between consecutive matches of the motif *M* in *S*. A sliding window approach through ***d***_*S*_ with window size *w* is used to obtain *n −* 1 *vectors of ranks*

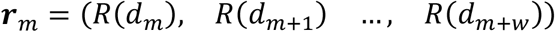

where *R*(*d*_*i*_) is the rank of *d*_*i*_ in the window. Given a *vector of ranks* ***r*** with window size *w*, we can describe a hash function

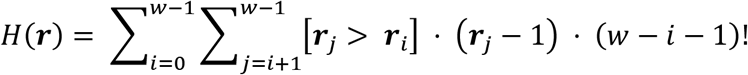

to represent the *vector of ranks* in a compressed form as a single positive integer. To account for reverse complementarity, a window [*d*_*m*_, *d*_*m*+*w*_] gets assigned a *sort space hash*

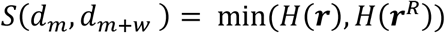

where ***r***^*R*^ is the *vector of ranks* as if the window was reversed

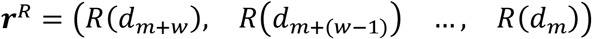

and ***r*** is the *vector of ranks* of the window. Then, in *permutation space*, these *sort space hashes* are mapped to each genome, which allows rapid lookup using a hash map data structure.

## Deconvolution

Because of sequence complementarity between related genome sequences, multiple genomes are expected to share *sort space hashes. Voyager* quantifies these overlaps in a square matrix. Given *k* indexed genomes [*g*_1_, *g*_*k*_], the *overlap matrix*

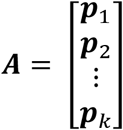

contains *k* row vectors ***p***, where

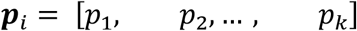

gives the proportion of expected hits in *g*_1_ through *g*_*k*_, given genome *g*_*i*_. Firstly, this information is used to deconvolute the raw obtained hits for sequence complementarity. Given the *overlap matrix* ***A*** and the *vector of observed hits* ***b***, either ***b*** ∈ *C*(***A***) *or* ***b*** ∉ *C*(***A***) where *C*(***A***) is the column space of the matrix ***A***. In other words, ***b*** may exist outside of the column space of ***A***. If ***b*** ∉ *C*(***A***), ***Ax*** = ***b*** has no solution for the *vector of deconvoluted hits* ***x***, because in this case, no linear combination of the column vectors of ***A*** can equal ***b***. However, the deconvoluted vector ***x*** ^∗^ can be approximated by minimizing the Euclidian norm of ***Ax*** ^∗^ *to* ***b***, *‖****Ax*** ^∗^ *−* ***b****‖* using the Householder’s transformations method for solving the NNLS problem as described by Lawson & Hanson^10^.

### Statistical significance estimation

*Voyager* also uses this *overlap matrix* to build a likelihood model for each of the genomes in the database, as well as a null hypothesis model. The null hypothesis model assumes that the proportions of observed hits are random and thus are distributed according to the respective proportion of each genome in *permutation space*. The alternative model assumes that the observed hits are distributed according to ***p***_*i*_. *Voyager* computes the log likelihood of such a model *θ* given a *vector of hits X* as

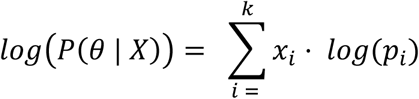

Then the probability of the genome can be derived from the likelihood of the null hypothesis model and the genome model according to

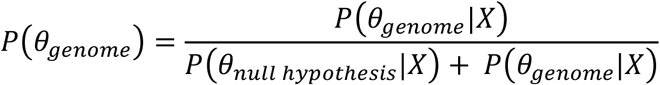

The corresponding p-value is computed using the chi^2^ cumulative distribution function chisq_cdf, and the likelihood ratio *LR* as follows: p-value ∼ 1 *−* chisq_cdf(*LR*; *DDOF*), where

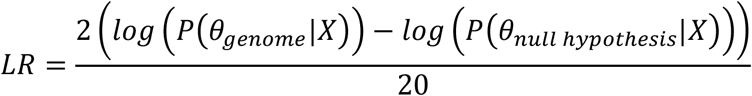

where 20 is an empirical scaling factor, and

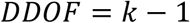

The chisq_cdf is computed using the lower incomplete gamma series expansion method as

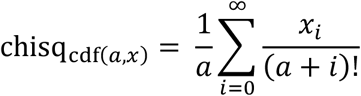

until convergence or the iteration threshold is reached.

### Reporting and Visualization

*Voyager* allows for taxonomic classification in real-time as the read data becomes available during the sequencing run. For monitoring the progress, *Voyager* implements a Graphical User Interface (GUI) written in Java, the *Voyager Monitor*, which receives ongoing updates from *Voyager* in JSON format via a network connection and displays the current results (Figure 8). The overview panel (left) reports the fraction of hits by genome either before or after deconvolution and the distribution of read lengths in the sample and lengths of mapped reads. The genome panel (right) shows details for each genome, the read length distribution, both absolute (light blue) and scaled on a per genome basis (gray), as well as the spatial distribution of hits throughout the genome.

**Figure 8:**
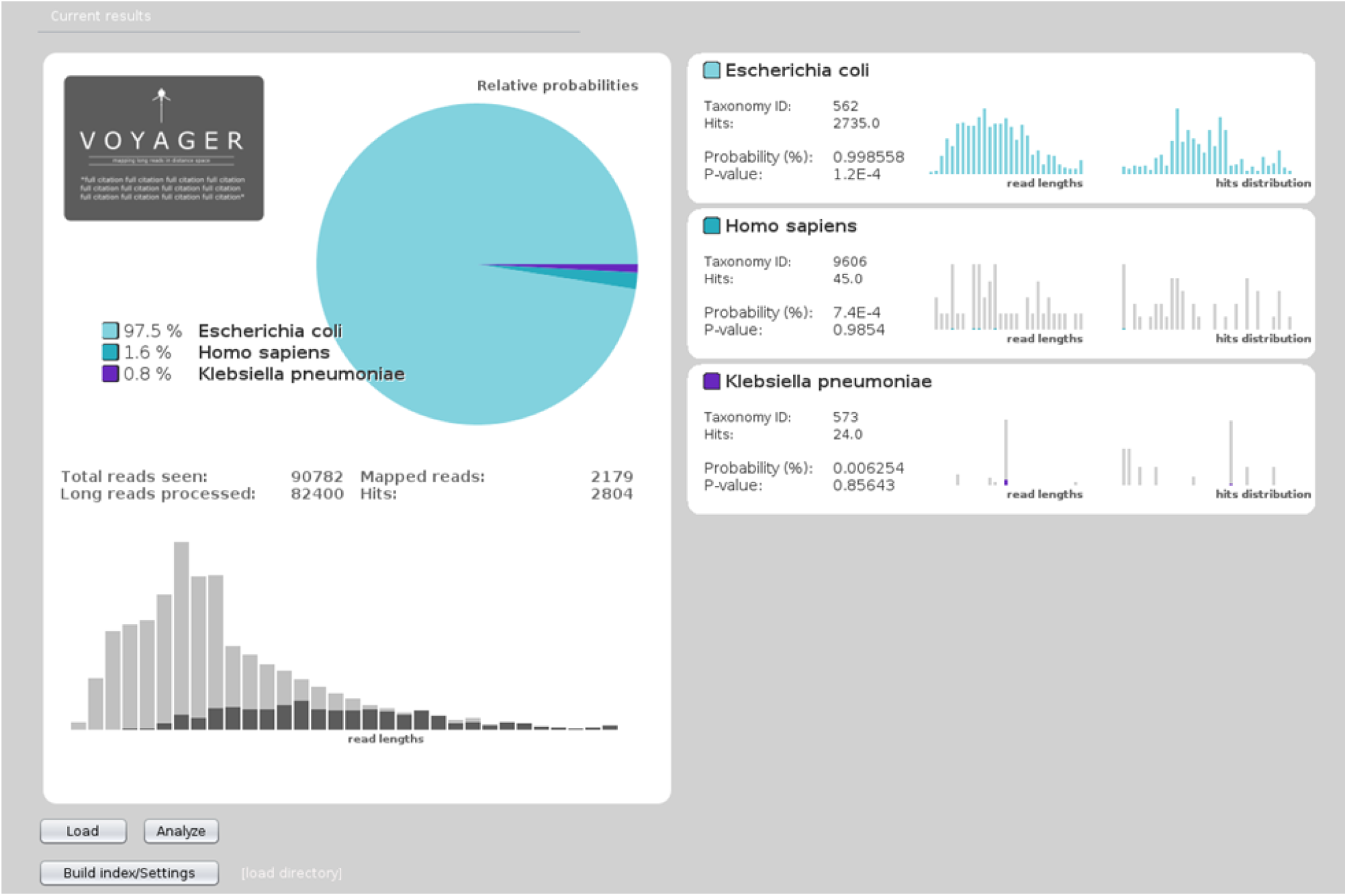
Screenshot of *Voyager Monitor*. The current taxonomic assessment is shown and updated while *Voyager* processes the data. Shown are: (a) a pie chart of the taxonomic distribution based on either the hits or the deconvoluted hits; (b) the distribution of read length, overlaid by the distribution of mapped reads (not to scale for better visual readability); and (c) a list of species detected, with the NCBI taxonomy ID, and distributions of read lengths and how they cover the respective genomes.

## Software Availability

*Voyager* is open-source software distributed under the LGPL and is available from https://bitbucket.org/sverre-phd-work/voyager/src/master/

## Glossary

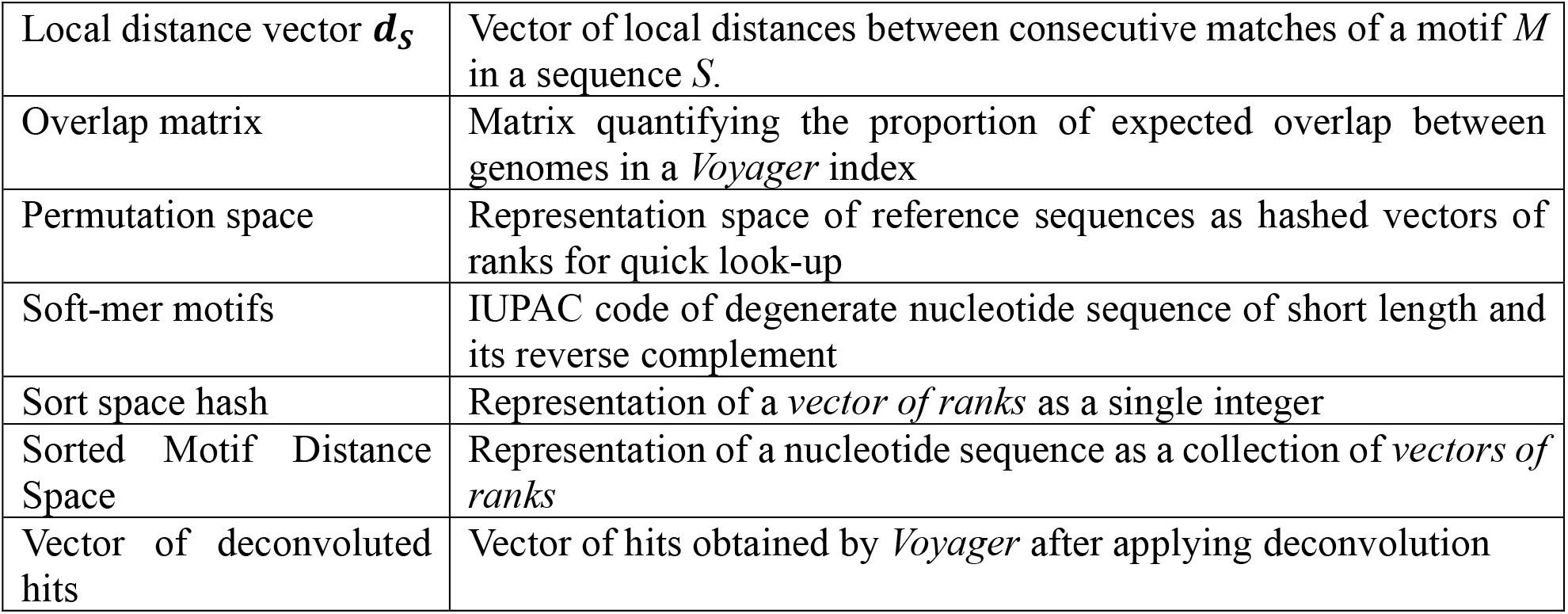
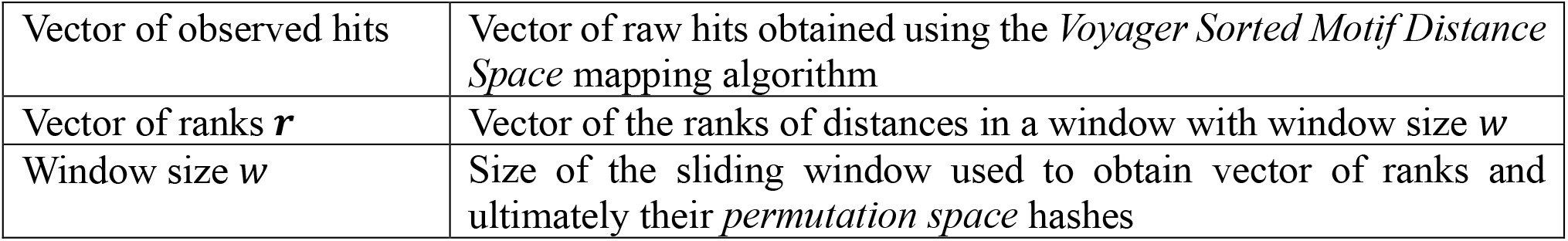

## Author information

### Authors and Affiliations

Department of Biotechnology, Inland Norway University of Applied Sciences, Holsetgata 22, 2317, Hamar, Norway

Sverre Branders, Manfred G. Grabherr & Rafi Ahmad

Institute of Clinical Medicine, Faculty of Health Sciences, UiT - The Arctic University of Norway, Hansine Hansens veg 18, 9019, Tromsø, Norway

Rafi Ahmad

## Contributions

RA, SB, and MGG designed the research. SB and MGG designed and implemented the software. SB performed the experiments and created the figures with input from RA and MGG. RA selected the test datasets. SB designed the user interface. SB, MGG, and RA wrote the manuscript.

